# Copy number networks to guide combinatorial therapy for cancer and other disorders

**DOI:** 10.1101/005942

**Authors:** Andy Lin, Desmond J. Smith

## Abstract

The dwindling drug pipeline is driving increased interest in the use of genome datasets to inform drug treatment. In particular, networks based on transcript data and protein-protein interactions have been used to design therapies that employ drug combinations. But there has been less focus on employing human genetic interaction networks constructed from copy number alterations (CNAs). These networks can be charted with sensitivity and precision by seeking gene pairs that tend to be amplified and/or deleted in tandem, even when they are located at a distance on the genome. Our experience with radiation hybrid (RH) panels, a library of cell clones that have been used for genetic mapping, have shown this tool can pinpoint statistically significant patterns of co-inherited gene pairs. In fact, we were able to identify gene pairs specifically associated with the mechanism of cell survival at single gene resolution. The strategy of seeking correlated CNAs can also be used to map survival networks for cancer. Although the cancer networks have lower resolution, the RH network can be leveraged to provide single gene specificity in the tumor networks. In a survival network for glioblastoma possessing single gene resolution, we found that the epidermal growth factor receptor (EGFR) oncogene interacted with 46 genes. Of these genes, ten (22%) happened to be targets for existing drugs. Here, we briefly review the previous use of molecular networks to design novel therapies. We then highlight the potential of using correlated CNAs to guide combinatorial drug treatment in common medical conditions. We focus on therapeutic opportunities in cancer, but also offer examples from autoimmune disorders and atherosclerosis.

## Introduction

New drug discovery is increasingly confronted by the obstacles of rising costs and diminishing success rates. To overcome these impediments, a number of strategies that employ genome and network data have been developed. These approaches can be used to reposition drugs outside their usual therapeutic domain or to design novel drug combinations. The networks that have been employed in these efforts typically consist of transcriptional co-expression networks or protein-protein interaction networks. However, these networks can be noisy and have many false-positives and false-negative observations. They also suffer from bias. Less effort has been devoted to constructing mammalian networks at the level of the gene. Here, we focus on genetic interactions in mammalian cells identified from correlated patterns of unlinked copy number alterations (CNAs). These networks represent an opportunity for the design of novel treatments and are particularly relevant to antiproliferative therapies for diseases such as cancer and autoimmunity, but can also be used for other disorders.

## A diminishing drug pipeline

A variety of therapeutic approaches are available for common disorders. Small organic molecules continue to be the mainstay of medical therapies, though prominent niche roles are progressively being taken by macromolecules, such as interfering RNA, gene therapy and therapeutic antibodies. Regardless of modality, it is increasingly difficult to gain approval for new drugs, leading to blocked pipelines for novel treatment strategies (Csermely et al., 2013; Gupta et al., 2013; Pujol et al., 2010; Zou et al., 2013). New drugs can fail at multiple steps in the testing process, often because of unexpected safety or toxicological concerns. Another daunting factor that discourages investment in new drugs are the enormous costs of development, which represent a high-stakes gamble even for a large company. In fact Eroom’s Law (Moore’s Law backwards), observes that the number of therapies developed per research dollar has halved every nine years for decades (Scannell et al., 2012; Wobbe, 2008).

## Eukaryotic molecular interaction networks

A variety of strategies are available to chart molecular networks. Protein-protein interactions can be identified using yeast two hybrid mapping or co-immunoprecipitation of protein complexes followed by mass spectrometry (Geva and Sharan, 2011; Giot et al., 2003; Venkatesan et al., 2009; Vidal et al., 2011; Yu et al., 2008). Genetic interaction networks have been dissected in yeast and worms. These networks have been mapped by seeking epistatic and synergistic interactions between alleles constructed by homologous recombination or RNA interference (RNAi) (Costanzo et al., 2010; Lehner et al., 2006). Interaction maps can also be charted by identifying groups of genes with correlated expression patterns (Zhang and Horvath, 2005). These groups of correlated genes are known as co-expression modules and employ undirected interactions. Directed networks can be mapped by relating the marker status of regulatory genes to the expression levels of target genes (Ahn et al., 2009).

A major problem with the available networks, particularly protein networks, is that they are noisy and have an appreciable number of false positive and false negative observations (Bruckner et al., 2009; Mackay et al., 2007). Because the networks are incomplete, they also suffer from bias, with preferential attention given to genes and proteins that have a high profile in the literature (Coulomb et al., 2005).

## Using genome data to replenish the drug pipeline

To replenish the diminishing flow of approved drugs, there has been growing interest in using the available variety of genomic and network data to design new drug therapies and minimize side effects. For example, one recent study evaluated transcript profiling data combined from many studies to identify CD44 gene expression as strongly correlated with type II diabetes mellitus (Kodama et al., 2012). Introducing CD44 deficiency into mouse models of diabetes blunted the effects of the disorder, suggesting that targeting this molecule will have useful therapeutic effects.

In addition to identifying new drug targets, molecular networks can be used to redeploy drugs approved for other disorders (Gupta et al., 2013; Zou et al., 2013). This approach is called drug repositioning or repurposing. One strategy employs transcript profiles in drug exposed cells, or other multidimensional readouts, to construct networks of drug-drug similarities. Modules of interconnected drugs can predict compounds that will have efficacy in settings where the drugs are not usually employed (Gottlieb et al., 2011; Iorio et al., 2010; Iskar et al., 2013). Other strategies for drug repositioning have used protein-protein interactions common to different drugs, constructing personalized drug networks from genome-wide association studies, and using drug side effects to point out novel therapeutic areas (Csermely et al., 2013; Pujol et al., 2010).

## The small world properties of networks facilitate combinatorial therapies

Biological networks typically display “small world” properties, whereby any two genes are separated by only a small number of links (Watts and Strogatz, 1998). If each gene interacts with 30 others, a gene will connect with 900 genes in two steps (i.e. 30^2^) and with all genes in the genome within three steps (30^3^ > 20,000 genes). In fact, the average path length in biological networks (the number of links between any two genes) varies between roughly 2 to 4 interactions. Thus, nearly all genes are linked within a short number of steps to all other genes (Albert, 2005; Albert and Barabási, 2002; Tsaparas et al., 2006; Vidal et al., 2011; Xu et al., 2011; Zou et al., 2012). There are nearly 3,000 Food and Drug Administration (FDA) approved drugs (http://www.fda.gov). After accounting for overlapping targets, it is estimated that these drugs affect approximately 1,000 different gene products (Overington et al., 2006). Approximately 1 in 20 genes is therefore a target for an FDA approved drug. Thus, within one step, each gene will interact with one to two drug targets, and within two steps, 45 targets.

The close connectivity of biological networks implies that they can be used to direct therapies towards disease genes and pathways by using approved drugs in single, double and triple combinations. In this way, causative genes can be targeted, even if there is no currently available treatment. Network guided combinatorial therapy can employ either repositioned drugs or drugs in their usual disease context. Despite the relatively small number of FDA approved compounds, network guided combinatorial strategies will permit a general, effective and accessible approach towards therapy (Kwong et al., 2012; Lin and Smith, 2011; Nijman and Friend, 2013; Pujol et al., 2010; Yang et al., 2010; Zou et al., 2013). Compared to the use of individual therapies, drug combinations exhibit greater efficacy with fewer side effects and decreased toxicity (Sun et al., 2013).

By restricting the search space for drug combinations, the network guided strategy offers a convenient approach to facilitate the development of drugs mixtures that are redeployed for diseases other than their original design. The use of already approved/developed drugs in network-guided combination treatments will speed therapy development and diminish the need for preclinical testing. Many orphan diseases, in particular, have no available drugs (Sardana et al., 2011). Network guided combinatorial therapy may provide treatment options for these disorders.

Further, the small world properties of biological networks may explain the common phenomenon in which unexpected therapeutic effects are obtained from drugs employed in diseases other than that for which originally designed. For example, thalidomide was initially developed as a sedative but is now used to treat cancer (Sissung et al., 2009).

## Molecular networks can be used to guide drug combinations

The small world properties of biological networks have been used with advantage to design combination therapies for a number of disorders, including cancer, diabetes, neurodegenerative disorders and infectious disease (Csermely et al., 2013; Kwong et al., 2012; Nijman and Friend, 2013; Pujol et al., 2010; Yang et al., 2010; Zou et al., 2013). A recent study used highly time resolved quantitation of transcript profiles and cell based phenotypes to show that EGFR inhibition reactivated apoptotic networks in breast cancer cells (Lee et al., 2012). These apoptotic pathways left the malignant cells susceptible to subsequent treatment with genotoxic drugs. Another investigation examined already employed therapeutic drug combinations and merged these data with known drug-target interactions and protein–protein interactions (Zou et al., 2012). The integrated data could be used to successfully predict new drug combinations.

A further approach employed an algorithm that incorporated previously reported drug-drug interactions to predict new interactions (Guimera and Sales-Pardo, 2013). Stochastic block models that used the notion of group-dependent interactions were employed to infer networks in which the interaction between any drug pair was predicted by the group in which the pair resides. Another study increased the efficiency of screens for drugs with synergistic interactions by combining pre-existing data from a small sample of empirically determined interacting drug pairs with other data, such as protein-protein interactions (Gerlee et al., 2013). A matrix algebraic technique based on cyclical projections onto convex sets significantly improved the rate of drug synergy discovery compared to traditional screens.

## Copy number alterations as a disease driver

For cancer, in particular, it is well established that amplification or deletion of genes plays a causative role. Amplification of the c-Myc gene and epidermal growth factor receptor (EGFR) genes, for example, have been strongly implicated in non-small cell lung cancer (NSCLC) (Sos et al., 2009) as well as a variety of other cancers (Beroukhim et al., 2010). Systematic surveys of DNA copy number alterations (CNAs), either deletions or amplifications, has linked cancer causation to an array of genes, either oncogenes or tumor suppressor genes. Genomic techniques can point to mechanisms by which the CNAs drive proliferation. In glioblastoma, CNAs have been connected to altered gene expression, which in turn has been related to survival (Jornsten et al., 2011). However, individual oncogenes have generally been studied in isolation. Co-inheritance patterns for pairs of amplified and deleted genes, particularly those distant from each other in the genome, have been subjected to more limited scrutiny.

## Using correlated copy number alterations to construct survival networks

Recent studies have sought genetic interaction networks for cancer by seeking correlated patterns of unlinked CNAs. Genetic survival networks identified using correlated CNAs have been found in glioma cells (Bredel et al., 2009; Rapaport and Leslie, 2010) and ovarian cancer cells (Gorringe et al., 2010). Correlated patterns of CNAs in cancer that span entire chromosome arms have also been identified (Kim et al., 2013). However, the chromosome arm network highlights a problem of charting CNA based interactions in cancer, which is that amplifications and deletions are not distributed randomly over the genome. Rather CNAs are flanked by hot spots for DNA rearrangements and can incorporate many genes (Beroukhim et al., 2010; Hsiao et al., 2013). This property can make identification of the causative gene pairs difficult.

## A pan-cancer CNA interaction network

In a recent study, the resolution of identified CNA interactions was improved by combining data from over 4,000 different cancers across 11 different varieties (Zack et al., 2013). For each cancer variety, there was a median of 74 consistent CNAs and a total of 770 CNA regions over all varieties. Pan-cancer CNAs were identified by combining the data from all cancers. The size of the significant CNAs decreased from 1.4 Mb in the individual cancers to 0.7 Mb in the pan-cancer CNAs, improving the resolution with which causative genes were mapped. However, by imposing the criterion that the CNAs were pan-cancer, the number of detectable events was diminished ∼5-fold. Further, most pan-cancer CNAs still harbored more than one gene, often more than 200. It was possible to construct a network by looking for correlated CNAs in the pan-cancer data. Not surprisingly, however, the size of the resulting network was small, with only 436 nodes.

## Mapping genetic survival networks using correlated CNAs in radiation hybrid cells

Our group has used radiation hybrid (RH) panels to map genetic circuits critical for cell survival. Radiation hybrid (RH) mapping was invented to determine the relative locations of genes within mammalian genomes (Cox et al., 1994; Goss and Harris, 1975). RH panels are constructed by lethally irradiating cells, causing the DNA to fragment into small pieces. The irradiated cells are then fused to living hamster cells, which incorporate the DNA fragments into their genomes. The resulting hybrid cells each contain extra copies of a random assortment of genes (∼25%), which are triploid rather than diploid. Genes in close proximity tend to be co-inherited in the RH clones, while genes far apart tend to be inherited independently. The small size of the DNA fragments affords the technique very high resolution, in fact, to within a single gene.

We showed that extra copies of distinct genes, *unlinked* triploid pairs, may enhance the survival of an RH cell (Lin et al., 2010). Because of the hardiness of the RH clones, statistically significant patterns of co-inherited genes pointed to the cell’s survival mechanism. Over 7.2 million statistically significant interactions were identified using the RH data, including genes that partner specifically with oncogenes. The RH network was mapped at single gene resolution (<150 kb) (Figure 1A) and the fact that the network was Gaussian rather than scale-free indicated that nearly all of the network has been charted. In fact, the RH survival network overlaps significantly with other protein-protein interaction networks, while being hundreds of times more comprehensive.

**Figure 1.**
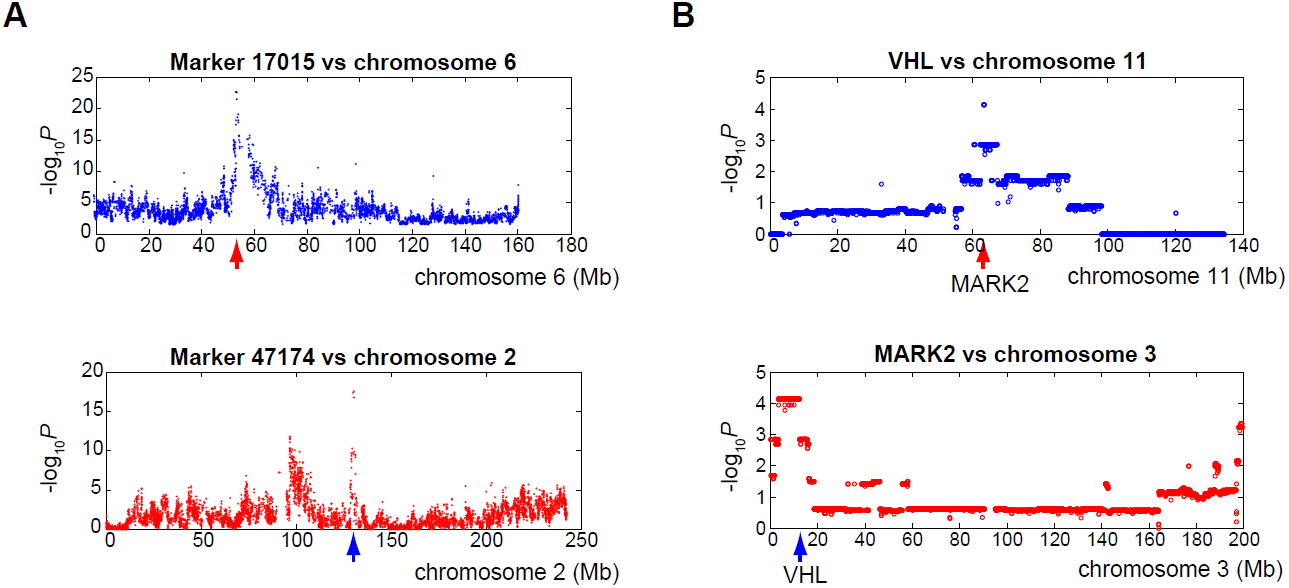
Genetic interactions in RH and GBM cells. (**A**) An interaction between a gene on chromosome 6 (red arrow) and a gene on chromosome 2 (blue arrow) in the RH network. The ordinate shows the significance value (-log_10_P) for co-retention. (**B**) An interaction between the MARK2 gene on chromosome 11 (red arrow) and the VHL gene on chromosome 3 (blue arrow) in the glioblastoma network.

## A survival network for glioblastoma multiforme at single gene resolution

We explored the existence of survival networks in cancer (Lin and Smith, 2011). Correlated patterns of copy number alterations (CNAs) for distant genes in glioblastoma multiforme (GBM) brain tumors were identified using the same method employed to construct the RH survival network. We analyzed public data on 301 glioblastoma multiforme brain tumors, which had been assessed for CNAs using array comparative genomic hybridization (aCGH) with 227,605 markers (The Cancer Genome Atlas (TCGA) Research Network, 2008). The tumors had a mean amplification length of 5.35 Mb and a mean deletion length of 5.87 Mb. A total of 11.2% genes were amplified in more than 5% of the glioblastomas and 0.9% deleted. Copy number variations found in the normal population were excluded.

Pairs of amplified genes in the tumors were identified that were separated by more than the corresponding upper limit of the amplification lengths in the genome. Pairs of distant genes both of which were deleted were also identified, or pairs of genes where one was amplified and the other deleted. We tested whether the amplification and/or deletion of the widely separated genes occurred simultaneously at a rate greater than by chance. A total of 436,302 interactions were found in the glioblastoma network at an FDR < 5%. An example of a gene interaction between the Von Hippel-Lindau (VHL) tumor suppressor gene and the MAP/Microtubule Affinity-Regulating Kinase 2 (MARK2) gene is shown in Figure 1B. Unlike the RH interaction peaks, the GBM interaction peaks have multiple plateaus, representing non-random breakpoints in the tumor DNA. This phenomenon decreases mapping resolution for interacting genes.

The glioblastoma and RH survival networks overlapped significantly (P = 3.7 × 10^−31^), validating the cancer network. We therefore exploited the high-resolution mapping of the RH data to obtain single gene specificity in the glioblastoma network. We identified overlapping interactions in the two networks to construct a combined network featuring 5,439 genes and 13,846 interactions (FDR < 5%). This network suggested novel approaches to the therapy of glioblastoma. An example using the epidermal growth factor receptor (EGFR) oncogene is discussed below.

## Using CNA networks to guide combination therapies

Although molecular networks have been employed to guide combinatorial therapies, limited attention has been devoted to the use of networks based on unlinked but correlated CNAs. In this article, we focus on gene interaction networks deduced from correlated CNAs in radiation hybrid (RH) cells and in cancer. The principal therapeutic opportunity using these networks is for disorders of cell proliferation, including cancer and autoimmunity. However, we also discuss atherosclerosis as a further example.

We illustrate three strategies by which CNA interaction networks can be used to design network guided combinatorial therapies; (**1**) Using subnetworks to identify multiple drug targets that interact with a single disease gene; (**2**) Using drugs to target multiple genes in a single disease pathway; and (**3**) Using drugs to target genes in parallel pathways converging on a single disease process. Drug/gene interactions In the examples were obtained from a number of databases, including DrugBank (http://www.drugbank.ca)(Knox et al., 2011) the Drug Gene Interaction Database (DGIdb; http://dgidb.genome.wustl.edu) (Griffith et al., 2013), GeneCards (Safran et al., 2010) (www.genecards.org), the Pharmacogenomics Knowledge Database (PharmGKB, http://www.pharmgkb.org) (Whirl-Carrillo et al., 2012) and the Therapeutics Targets Database (http://bidd.nus.edu.sg/group/ttd/ttd.asp) (Zhu et al., 2012). Other databases can also be employed (Csermely et al., 2013; Sun et al., 2013; Zou et al., 2012).

## Targeting multiple drugs to single disease genes in cancer

The c-Myc oncogene plays a major role in a wide variety of cancers (Wang et al., 2011). No approved compounds are available that specifically inhibit c-Myc, but a strategy that targets genes interacting with this gene product may be fruitful (Yang et al., 2010). In the RH survival network, 45 genes were linked with statistical significance (false discovery rate, FDR < 10^−4^) to c-Myc. Of the genes that interacted with c-Myc in the RH network, 12 (27%) happened to be specific targets for already existing drugs, though not necessarily for cancer treatment (Figure 2A). For example, the BMI1 polycomb ring finger oncogene product (PCGF4) is a subunit of an E3 ubiquitin ligase and is inhibited by the compound PRT4165 (Alchanati et al., 2009). Similarly, MAP2K5 (MEK5/ERK5) is a dual specificity protein kinase belonging to the MAP kinase kinase family and is inhibited by the compounds BIX02188 and BIX02189 (Tatake et al., 2008).

**Figure 2.**
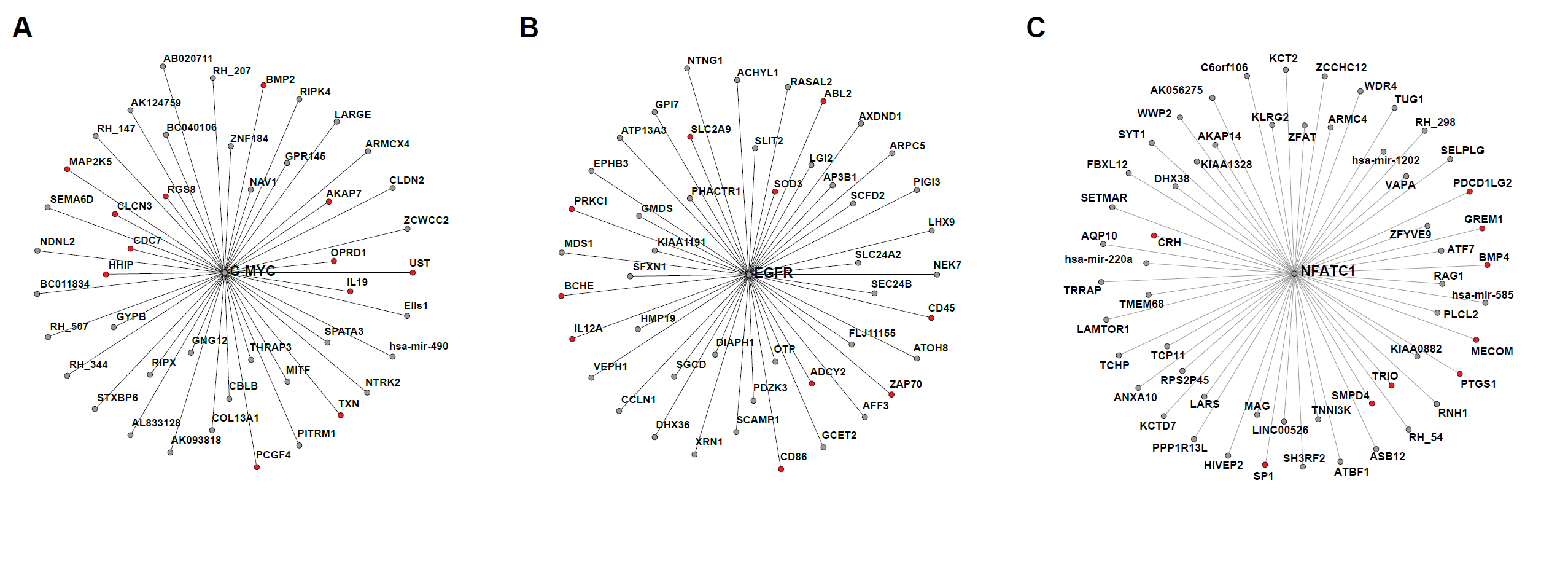
Using subnetworks to target individual node genes. (**A**) Subnetwork for c-Myc and all genes one edge away (FDR < 10^−4^). Genes in red are targets for existing drugs. (**B**) A subnetwork for the EGFR gene in the combined RH/glioblastoma network (FDR < 0.05). (**C**) Subnetwork for the T cell activation gene NFATc1 and all genes one edge away (FDR < 10^−4^).

The epidermal growth factor receptor (EGFR) oncogene is frequently activated in glioblastoma and other cancers. Medications that target the EGFR oncogene include the monoclonal antibody cetuximab (Erbitux) and the kinase inhibitors erlotinib (Tarceva) and gefitinib (Iressa) (Stinchcombe et al., 2010). Eventually, however, resistance to these treatments occurs (Dhomen et al., 2012).

A total of 46 genes were identified that interacted with EGFR in the combined glioblastoma/RH survival network (FDR < 0.05), of which 10 (22%) happened to be targets for existing drugs (Figure 2B). For example, butyrylcholinesterase (BCHE) is inhibited by donepezil, an anticholinesterase employed in treatment of Alzheimer’s disease (Anand and Singh, 2013). SLC2A9 is a high capacity urate transporter and is inhibited by the uricosuric agent benzbromarone which is used to treat gout (Caulfield et al., 2008; Doring et al., 2008; Vitart et al., 2008). These observations suggest that a flank attack strategy which strikes at both EGFR and its partner genes in the glioblastoma survival network may be an effective approach to treatment of these tumors. The genetic survival network for glioblastoma thus offered insights into the mechanisms of proliferation for this cancer and suggested new avenues for therapeutic intervention.

Patient-to-patient variations exist in disease networks. For instance, a variety of oncogenes are activated in different cancers (The Cancer Genome Atlas (TCGA) Research Network, 2008; Zack et al., 2013). Our strategy of using correlated CNAs to guide combination therapies can account for individual variations in disease networks, providing a foundation for personalized medicine.

## Targeting multiple drugs to a single disease gene in autoimmunity

Another example employing correlated CNA networks is provided by NFATc1. This gene plays a key role in T cell activation, an important cellular response in autoimmune disorders (Bartelt et al., 2009; Kannan et al., 2012; Smith-Garvin et al., 2009). In the RH survival network, 56 genes were linked with statistical significance (FDR < 10^−4^) to NFATc1 (Figure 2C). No approved compounds exist that specifically target NFATc1. However, of the genes that interact with NFATc1, 9 (16%) happen to be specific targets for already existing drugs. One unsurprising example is PTGS1 (cyclooxygenase 1). This enzyme is involved in prostaglandin synthesis and is a target for non-steroidal inflammatory drugs (NSAIDs) (Dinarello, 2010). Another plausible example is the MECOM oncoprotein, which is specifically degraded by arsenic trioxide (ATO), perhaps explaining the promise of this compound in treatment of autoimmune syndromes (Bobe et al., 2006; Shackelford et al., 2006). Other interacting genes and their cognate pharmaceuticals were more unexpected and have yet to be utilized to treat autoimmune conditions The enzyme SMPD4 (sphingomyelin phosphodiesterase 4) is inhibited by the compound GW4869 (Chipuk et al., 2012). Similarly, the TRIO gene encodes a rho guanine nucleotide exchange factor, which is specifically inhibited by the compound ITX3 (Bouquier et al., 2009).

## Targeting multiple genes in a single disease related pathway in cancer

The second strategy that employs CNA network information uses multiple drugs to perturb genes participating in a single pathogenic pathway. An example of one such pathway in the RH network which ends on the EGFR oncogene is shown in Figure 3. (Note that this subnetwork does not incorporate information from the glioblastoma CNA network and may be a more general network than e.g. Figure 2B.) A total of 22 genes interacted with the EGFR gene in the RH network (FDR < 10^−6^), of which seven (32%) happened to be targets for existing compounds (Figure 3). For example, R59022 inhibits diacylglycerol kinase β (DGKB) (Batista et al., 2005; Kamio et al., 2010) and RGB-286147 inhibits PFTAIRE protein kinase 1 (PFTK1) (Caligiuri et al., 2005).

**Figure 3.**
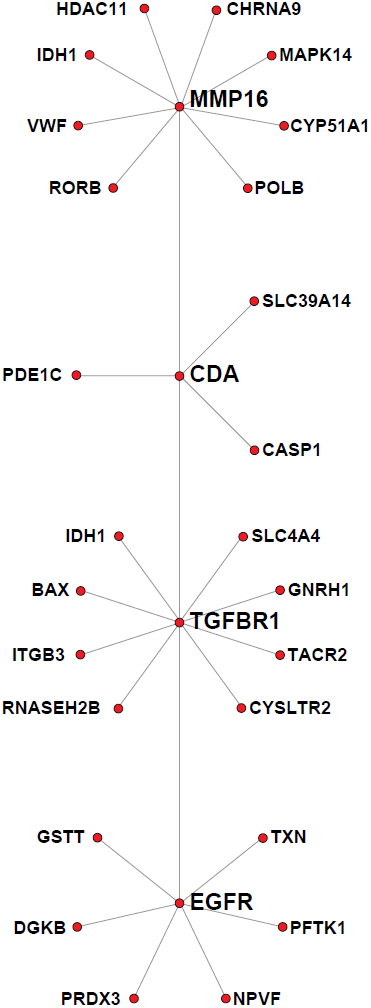
Targeting an individual pathway. A pathway leading to the EGFR oncogene in the RH network. Genes in red are targets for existing drugs. Genes that are non-drug targets are not shown. (FDRs for interacting genes: MMP16 < 10^−5^, CDA < 10^−4^, TGFB1 < 10^−5^, EGFR < 10^−6^.)

A trio of gene products that interacted with EGFR had antioxidant activity (Figure 3). Thioredoxin reductase (TXN) is inhibited by the gold compound, auranofin (Cox et al., 2008; Liu et al., 2012), which is also employed to treat autoimmune conditions such as rheumatoid arthritis. Peroxiredoxin 3 (PRDX3) is inhibited by thiostrepton, a thiazole antibiotic that shows activity against tumor cells (Newick et al., 2012). Glutathione-S-transferase (GSTT) is inhibited by α tocopherol, a form of vitamin E (Van Haaften et al., 2001), as well as by ellagic acid and curcumin, plant polyphenolic compounds (Hayeshi et al., 2007). There has been rising interest in inhibiting reduction/oxidation pathways for cancer treatment, since these pathways are required for cell proliferation (Kwok et al., 2008; Newick et al., 2012; Tew and Townsend, 2011). One mechanism by which these pathways might exert their therapeutic effects may be exemplified by interactions with oncogenes such as EGFR.

The transforming growth factor β receptor 1 gene (TGFBR1) also interacted with EGFR (Figure 3). TGFBR1 is a target for a number of kinase inhibitors, including SB525334 and SD-208 (Akhurst, 2006; Mohammad et al., 2011; Thomas et al., 2009). In addition, a total of 72 genes interacted with TGFBR1 (FDR < 10^−5^), of which 9 (13%) represented targets for available drugs. One of these genes was cysteinyl leukotriene receptor 2 (CYSLTR2), which is inhibited by available leukotriene inhibitors such as zafirlukast and zileuton. These compounds are used clinically as anti-inflammatory agents (Scow et al., 2007). Another gene that interacted with TGFBR1 was tachykinin receptor 2 (TACR2). Antagonists of this receptor include ibodutant and saredutant (Santicioli et al., 2013). TGFBR1 also interacted with cytidine deaminase (CDA), which in turn interacted with matrix metalloproteinase-16 (MMP16) (FDR < 10^−4^). CDA is inhibited by chemotherapeutic drugs such as tetrahydrouridine (Beumer et al., 2008) and zebularine (Lemaire et al., 2009). MMP16 is inhibited by marimastat (Wong et al., 2013). Both CDA (interactors FDR < 10^−4^) and MMP16 (interactors FDR < 10^−5^) were linked with a number of additional genes whose products can be antagonized by available drugs (Figure 3). The wide variety of available drugs that target the EGFR pathway suggest that combinations of these drugs might have therapeutic benefits in applications in which the EGFR gene is a key node driving oncologic proliferation.

## Targeting genes in parallel pathways converging on atherosclerosis

The third strategy that uses CNA networks to design new therapies is to direct drugs towards parallel disease pathways. As an example of this strategy, multiple pathways have been implicated in atherogenesis (Figure 4) (Lusis, 2012; Lusis et al., 2004). Apolipoprotein B (APOB) is the major protein constituent of low density lipoprotein (LDL) and elevated LDL concentrations are associated with increased atherosclerotic risk. Lipoprotein (a) (LPA) is also a lipoprotein that raises the risk of atherosclerosis through unknown mechanisms. The zinc fingers and homeoboxes 2 gene (ZHX2) and the Ox40 ligand (TNFSF4) have been implicated in atherosclerosis through genetic studies in mice and humans. There are no available drugs that directly affect any of these proteins. However, each of these atherogenic genes interact with between 5 to 9 genes (FDR < 10^−4^) that are targeted by available compounds. Some of these drugs are already employed as anti-atherogenic agents.

**Figure 4.**
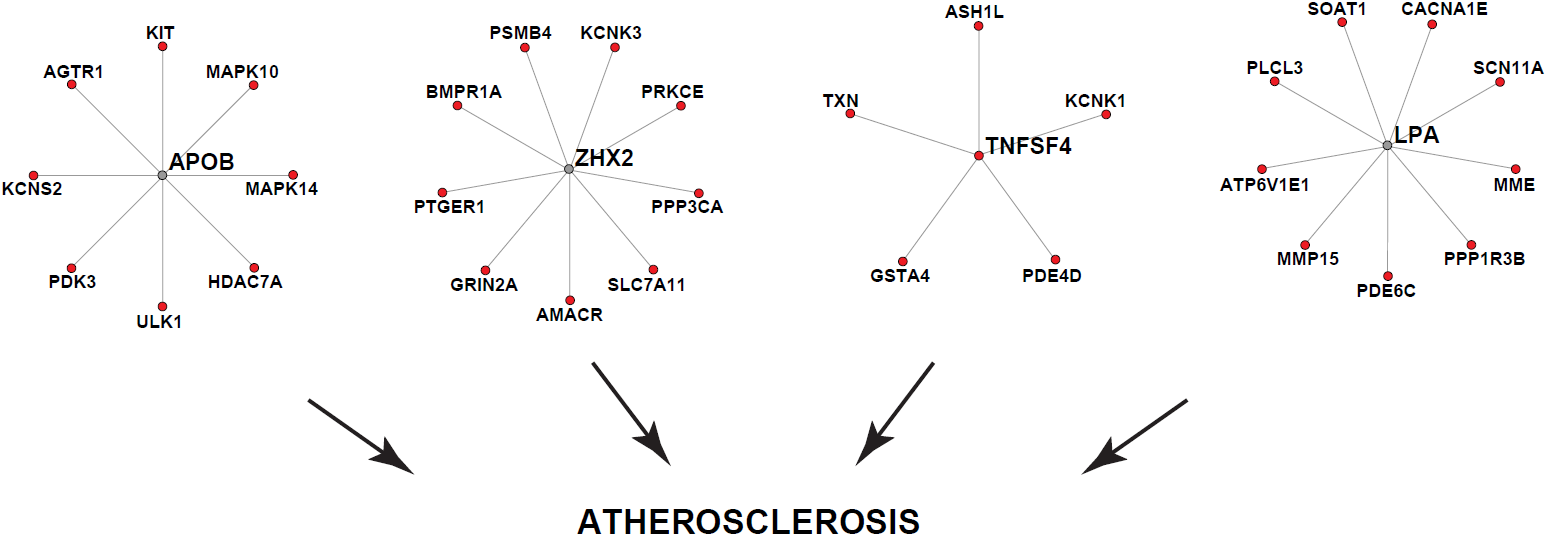
Targeting parallel pathways. Genes that conspire to promote atherogenesis in the RH network. Genes in red are targets for existing drugs. Genes that are non-drug targets are not shown. (FDRs for interacting genes < 10^−4^).

For example, ZHX2 interacts with the prostaglandin E receptor 1 gene (PTGER1). Non-steroidal anti-inflammatory drugs (NSAIDs), such as aspirin and naproxen, inhibit the cyclooxygenase enzymes which synthesize the prostaglandin ligands for this receptor. NSAIDs are widely used as prophylactic drugs to protect against atherosclerosis. The APOB gene interacts with the tyrosine kinases c-Kit (KIT) and MAPK14. KIT can be inhibited by kinase inhibitors such as imatinib and dasatinib (Ashman and Griffith, 2013). Similarly, phosphorylation of MAPK14 can be blocked using the kinase inhibitor sorafinib (Chapuy et al., 2011). The conjecture that kinase inhibitors may be beneficial in atherosclerosis is supported by recent studies (Grimminger et al., 2010; Hilgendorf et al., 2011).

The network connections of drug targets may explain their unexpected therapeutic effects. The angiotensin II receptor, type 1 (AGTR1) is significantly linked to APOB in the RH network (Figure 4). The angiotensin converting enzyme (ACE) inhibitors (e.g. enalapril), and the angiotensin receptor blockers (ARBs) (e.g. losartan) are effective in combating atherosclerosis (Patarroyo Aponte and Francis, 2012). The connection of AGRT1 with APOB might explain part of the efficacy of angiotensin pathway blocking agents as anti-atherosclerotic drugs, in addition to their role as antihypertensive agents.

## Using CNA networks to synergize drug combinations and minimize side effects

Drug combinations targeted to parallel or convergent pathways might allow the use of otherwise low efficacy drugs. One potential example of this synergistic strategy is provided by marimastat, which inhibits MMP16 in the RH pathway terminating on EGFR (Figure 3). Marimastat is not used clinically because of an unacceptable side effect profile (Wong et al., 2013). By combining marimastat at low concentrations with other drugs in a network guided strategy, it might be possible to maximize their common therapeutic effects, while minimizing their divergent adverse effects. However, accumulating side-effects will eventually set limits to polypharmacy. The optimal balance between therapeutic synergism and gathering side-effects will require empirical investigation.

Based on network data alone, it is not always possible to predict the direction of a drug effect. For example, APOB interacts with histone deacetylase 7A (HDAC7A) (Figure 4). HDAC7A is a class II HDAC, and is a target for inhibition by histone deacetylation inhibitors (HDIs). In fact, recent studies indicate that HDIs show promise in the therapy of atherosclerosis (Ordovas and Smith, 2010; Xu et al., 2012; Zhou et al., 2011). However, the HDI trichostatin A targets HDAC7A, but is proatherogenic in mouse models (Choi et al., 2005), underlining the necessity of experimental testing. Nevertheless, the strategy of CNA network guided combinatorial therapy promises to be a useful approach to advancing novel treatments for a wide variety of common and uncommon disorders.

